# New genetic sources for orange color in cucumber (*Cucumis sativus L.*) fruit flesh

**DOI:** 10.1101/685289

**Authors:** Brian M. Waters, HaeJin Kim, Keenan Amundsen

## Abstract

Most cucumber varieties have fruits with white flesh, which is devoid of ß-carotene and has a low concentration of total carotenoids. Carotenoids are important nutrients for humans and animals. Thus, developing cucumber varieties with orange flesh could provide a new nutritionally enhanced food source. Some cucumbers with yellow and orange flesh have been described, but there are others that have not been studied. Here, we used three cucumber PI lines, reported to produce colored fruits, from the USDA National Plant Germplasm System to generate three F_2_ populations. Fruits from the F_2_ populations with colored flesh (green, yellow, or orange) were pooled, and an equal number of fruits with white flesh were pooled. RNA was isolated from the pools and used for RNA sequencing to determine gene expression differences and to identify SNPs in each pool. The orange color of the cucumber fruits was confirmed to be due to ß-carotene. There were no clear expression patterns for genes of the carotenoid biosynthesis pathway that would suggest that their expression controlled the coloration of fruits. Mutations in carotenoid biosynthesis genes also did not explain the variation. However, we detected a SNP in the homolog of the *Or* gene that is responsible for ß-carotene accumulation in orange cauliflower. This genetic basis is different from that of previously studied orange cucumbers, but our results suggest that *Or* is not the only factor. These results provide the basis for future studies for breeding orange cucumbers for commercial or home garden production.

## Introduction

Cucumber (*Cucumis sativus* L.) is a popular and widely consumed fruit. Most cucumber varieties have green skin (exocarp) and white flesh (mesocarp and endocarp) (Che and Zhang, 2019). A number of genes that control exocarp color, which can also be orange, yellow, or cream-colored, have been described (Che and Zhang, 2019). Some cucumber varieties have colored flesh, such as yellow (Kooistra, 1971, Lu, H. W. *et al.*, 2015) or orange (Simon and Navazio, 1997), due to carotenoid accumulation (Navazio and Simon, 2001, Cuevas *et al.*, 2010), while the white flesh has no detectable ß-carotene and only trace amounts of total carotenoids (Kandlakunta *et al.*, 2008). Carotenoids are yellow, orange, or red compounds with various functions in plants (Cazzonelli and Pogson, 2010). These carotenoid compounds also have important nutritional value for humans and animals, such as ß-carotene serving as a provitamin A compound (Nisar *et al.*, 2015). Increasing the quantity of ß-carotene in cucumber could positively contribute to improved human nutrition and may provide a specialized horticultural product that is a colorful option for consumers. Colored cauliflower, carrots, and potatoes are popular with consumers. Orange cauliflower is caused by a mutation in a gene called *Orange* (*Or*) that encodes a DnaJ protein (Lu, S. *et al.*, 2006). The *or* mutation results in development of chromoplasts (Lu *et al.*, 2006) where ß-carotene accumulates (Li, L. and Yuan, 2013). SNPs in the homolog of this gene in melon, *CmOr*, can determine whether fruits are orange or green (Tzuri *et al.*, 2015).

Another way that carotenoid levels can be controlled is through the carotenoid biosynthesis pathway. Expression levels of a plant phytoene synthase (*PSY*) and a phytoene desaturase (*PDS*), the first enzymes in the carotenoid synthesis pathway, are key contributors to ß-carotene levels in plants (Cazzonelli and Pogson, 2010). Additionally, decreased expression or mutation of proteins downstream of ß-carotene, such as ß-carotene hydroxylase (BCH), can also affect ß-carotene accumulation by disrupting its conversion to downstream compounds (Qi *et al.*, 2013, Diretto *et al.*, 2007).

Some cucumber germplasm that produces fruits with colored flesh have been studied. The line PI 200815 produces fruits with yellow flesh, but the genetic basis has not been determined (Kooistra, 1971, Lu *et al.*, 2015). Orange colored germplasm has been identified and studied using the Xishuangbanna (XIS) cucumber from China (Simon and Navazio, 1997, Navazio and Simon, 2001, Cuevas *et al.*, 2010, Bo *et al.*, 2012). The orange coloration is due to ß-carotene (Cuevas *et al.*, 2010), and a mutation in the *BCH1* gene was identified as the cause of orange coloration in a GWAS study (Qi *et al.*, 2013).

Other than XIS and PI 200815, sources of colored flesh cucumber germplasm have not been studied, although there are some PI lines in the USDA National Plant Germplasm System that are described as producing fruits with colored flesh. The objectives of this study were to observe molecular differences in white and colored pools of cucumber fruits in F_2_ populations derived from PI 200815, PI 163217, and PI 606066. Our approach was to perform RNA sequencing in these pools and observe differentially expressed genes (DEGs) and SNP differences between the pools, with an emphasis on genes of the carotenoid biosynthesis pathway and DnaJ chaperones. Our results indicate that the underlying basis for colored flesh differs from that of the XIS cucumber, and there may be multiple genes that will need to be identified to fully understand control of flesh color. These results are an important step for developing new sources of orange flesh genes for cucumber breeding.

## Materials and Methods

### Plant materials

Seeds of cucumber lines PI 606066, PI 163217, and PI 200815 were obtained through the USDA National Plant Germplasm System from the North Central Regional Plant Introduction Station in Ames, Iowa. PI 606066 was collected from Madya Pradesh, India during the rainy season, and is described as having “orange flesh like *C. melo*” in the U.S. National Plant Germplasm System. PI 163217 was collected from Punjab, Pakistan, and is described as “flesh bright salmon”. PI 200815 was collected from Mandalay, Myanmar, and is described as having “orange flesh”.

Plants of each line were grown in a field and crossed with each other line. F_1_ plants were grown in a greenhouse and self-pollinated to produce F_2_ populations. On May 18, 2018, 25 plants of each population were transplanted into a field site on the University of Nebraska campus, and were allowed to be open-pollinated. Flowering began around the third week of June, and two representative fruits were harvested from each plant on July 23, 2018. Some plants had died, had been eaten by deer, or did not produce fruits. Fruit weight was recorded, as well as exocarp color and flesh color. Fruits without color (white flesh) and fruits with coloration (green, yellow, or orange) in the mesocarp and endocarp were selected for two pools of 17 fruits each. A cork borer was used to sample a plug of each fruit, which included exocarp, mesocarp, and endocarp, but not seeds or placental material. The plugs were frozen at -80 C until being ground together in a large mortar in liquid nitrogen. Four samples from each pool were used for RNA isolation using the Plant RNeasy kit (Qiagen).

### RNA-seq and SNP analysis

The three best quality RNA samples were subjected to RNA sequencing using the 2 × 75 bp High-Output Kit on an Illumina NextSeq 500 (University of Nebraska Medical Center DNA Sequencing Core Facility, Omaha, NE). Raw sequencing reads were deposited to the short read archive National Center for Biotechnology Information database under the BioProject PRJNA546604 (https://www.ncbi.nlm.nih.gov/bioproject/?term=PRJNA546604). Trimmomatic 0.36 (Bolger *et al.*, 2014) was used to remove low quality reads and reads contaminated with primers and adaptor sequences from the Illumina sequencing kit. During the trimming process, reads were also trimmed to a uniform 50 bp length. The Gy14 v2 transcriptome was obtained from CuGenDB (Zheng *et al.*, 2019). Forward sequencing reads from each pool were separately mapped to the Gy14 transcriptome using Bowtie 2.3 (Langmead and Salzberg, 2012). A read count data matrix was constructed and the Bioconductor edgeR package (Robinson *et al.*, 2010) was used to infer differential gene expression between color and white pools. Differentially expressed genes were identified as described previously (Waters *et al.*, 2014) from transcripts having at least 100 mapped reads and an edgeR false discovery rate < 0.1.

Single nucleotide polymorphisms and relatively short insertions and deletions relative to the Gy14 transcriptome were identified using the default settings of Samtools 0.1 mpileup (Li, H. *et al.*, 2009). Variants conserved among reps of one condition (color or white), but that differed from the other condition, were identified.

### Carotenoid analysis

Cucumber flesh samples from one orange-fleshed, one white-fleshed, and one green-fleshed fruit were lyophilized. About 100 mg of lyophilized sample was put into a screw-cap glass tube (13 × 100 mm, 9 mL) and 2 ml methanol: dimethylchloroether (9:1) was added. To extract carotenoids, the samples were ground with a homogenizer, then incubated at room temperature for 30 min. The glass tubes were centrifuged, and 1 ml solution was used for HPLC analysis. All samples were shielded from light during all processes. HPLC analysis was performed on an Agilent 12000 series system, fitted with a ProntoSIL 200-5 C30 column. The mobile phase was methanol: tert-butyl methyl ether (80 : 20, v/v) at 1 mL/min flow rate. Detection wavelength was 455 nm.

## Results

### Fruit size and color variation

Fruit size (mass) varied in each F_2_ population (Fig. 1). The two populations that were generated with the PI 200815 parent produced smaller fruits than the third population. The PI 200815 X PI 163217 population fruits ranged from 511 g to 1740 g, with an average mass of 998 g. The PI 606066 X PI 200815 population fruits ranged from 333 g to 1470 g, with an average mass of 851 g. The PI 606066 X PI 163217 population fruits ranged from 381 g to 2816 g, with an average mass of 1495 g. Fruit skin (exocarp) color also varied greatly (Fig. 2A, Table 1, Table S1). When combining observations from all three populations, green was the most frequent skin color, followed by orange, then yellow. However, skin color proportions differed between the populations.

**Table 1.**
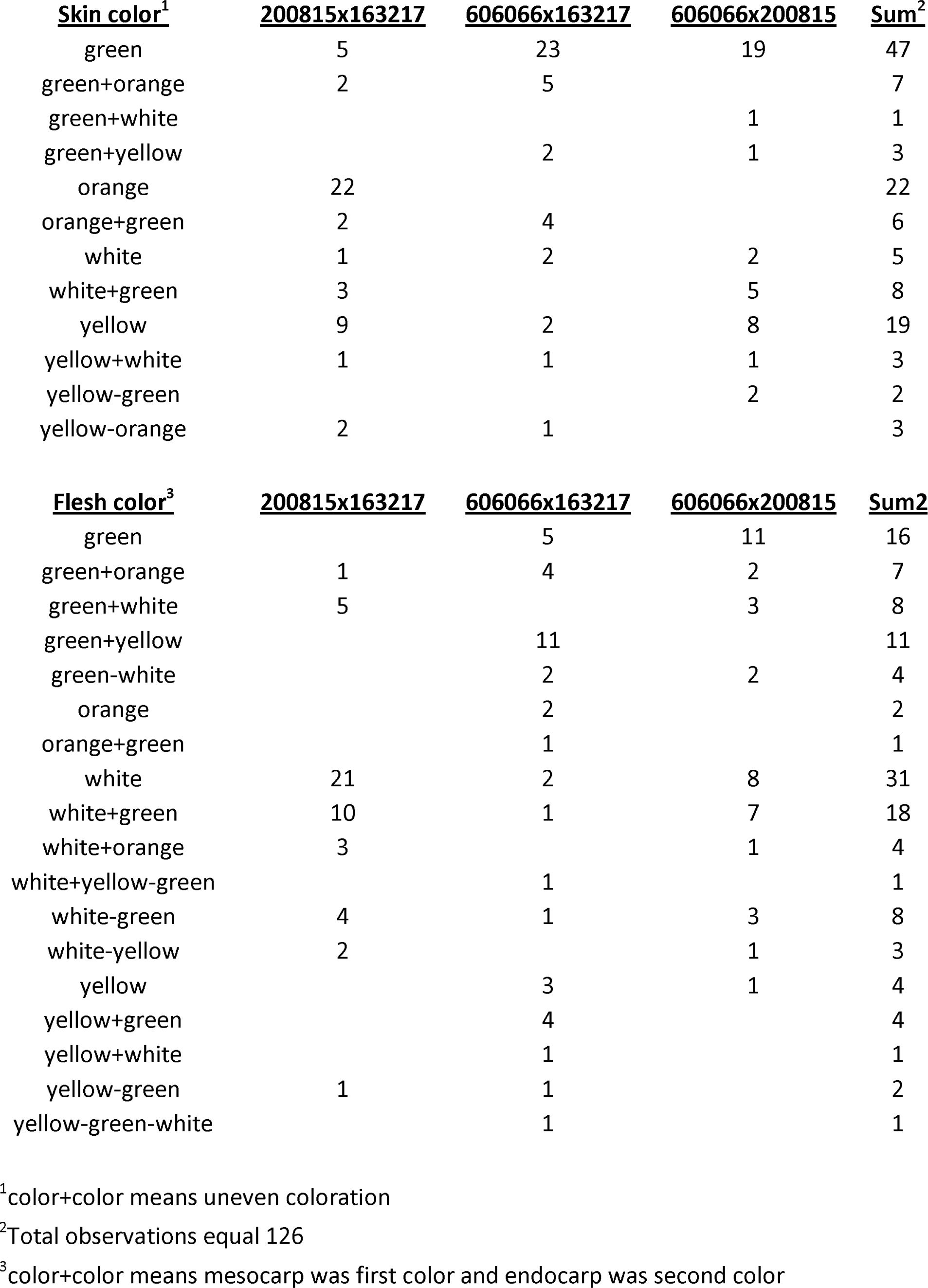
Summary of skin and flesh colors in fruits of three cucumber F_2_ populations.

**Figure 1.**
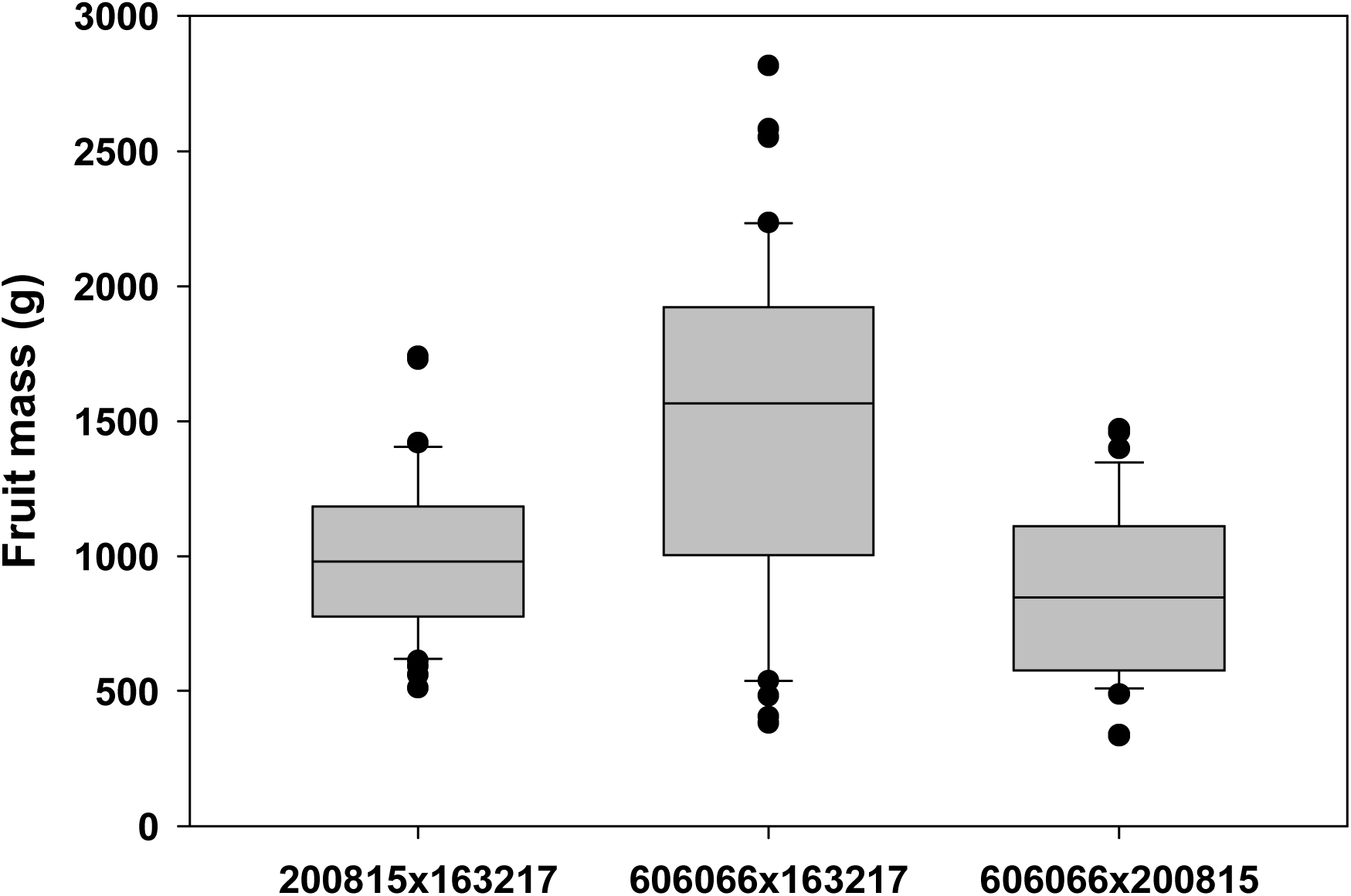
Cucumber fruit size (mass) of fruits from each of three F_2_ populations used in this study. Two fruits were collected from each plant. Numbers indicate each PI line parent.

**Figure 2.**
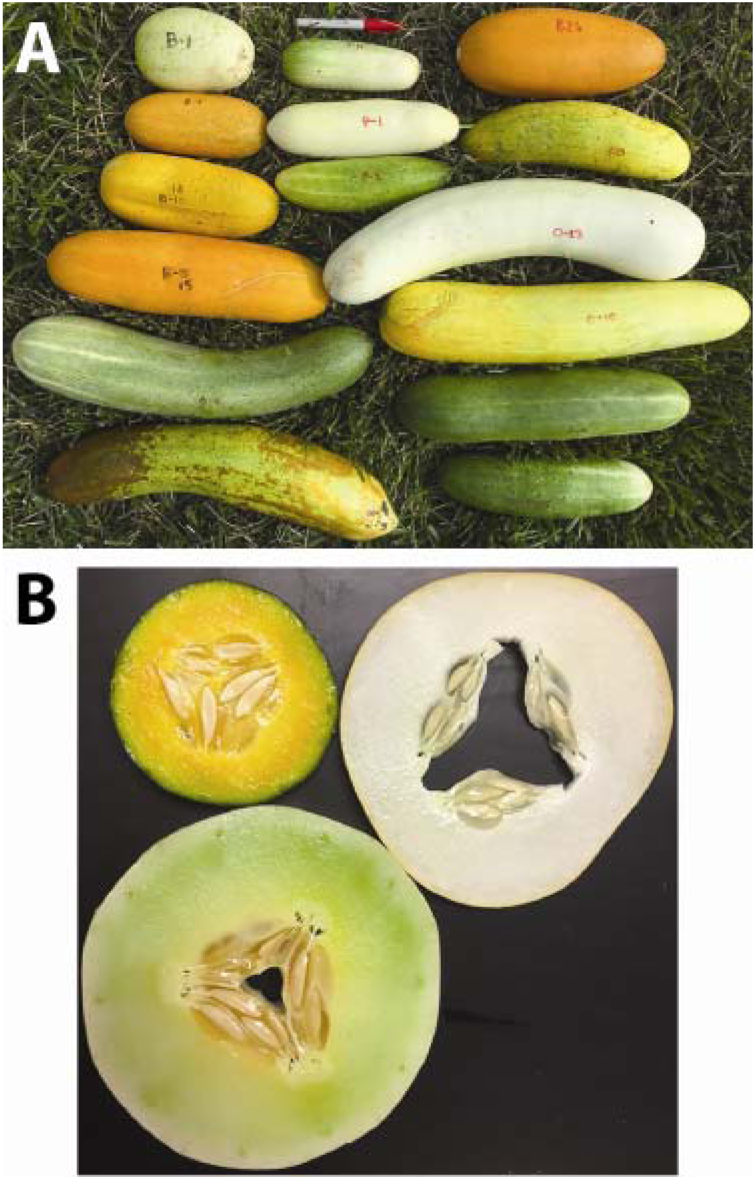
Diversity of cucumber fruit size, shape, and color. A, Photograph of several fruits from each F_2_ population. B, Cross-section photograph of orange, green, and white fruits used for HPLC analysis of beta-carotene.

Fruit flesh (mesocarp and/or endocarp) color also varied within each population (Table 1, Table S1). Combining observations from all three populations, the most frequent flesh color in the F_2_ fruits was white, followed by white with some coloration in the endocarp. Occasionally there were fruits that had green or yellow flesh, and two fruits from the same plant from the PI 606066 X PI 163217 population had orange flesh. We found 17 fruits that had flesh with substantial coloration, and pooled these fruits (Table S1, Table 2) for RNA sequencing, along with a separate pool of 17 white fruits.

**Table 2.**
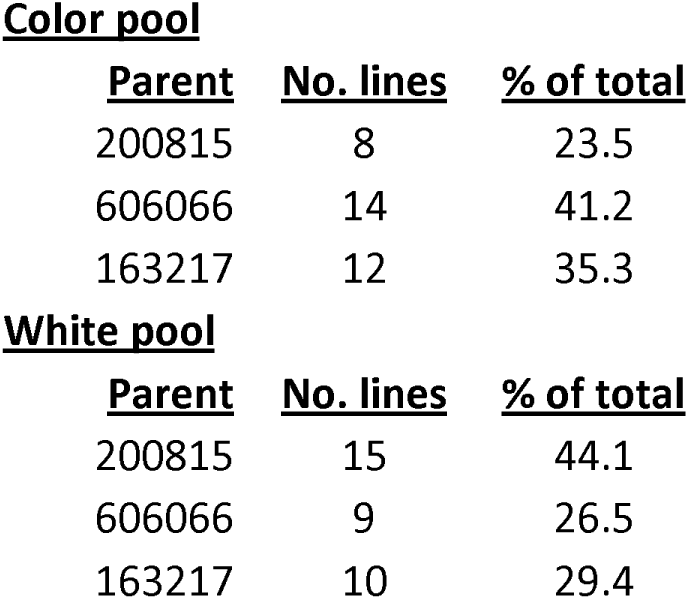
Contribution of each parent to color and white flesh pools.

### Carotenoid pathway and DnaJ chaperone gene expression

From the RNA sequences from the color and white pools, we were able to determine differential gene expression between the pools. 1889 genes had higher expression in the color pool, and 1292 genes had higher expression in the white pool (Supplementary Table S2). From among these DEGs, we focused on expression of genes in the carotenoid synthesis pathway (Table 3). Expression of some genes that are upstream of ß-carotene, such as PSY and carotene cis-trans isomerase (CRTISO) had slightly higher expression in the color pool, yet some other genes that are upstream of ß-carotene, such as zeta-carotene isomerase (ZISO) and zeta-carotene desaturase (ZDS), had higher expression in the white pool. Likewise, expression of some genes downstream of ß-carotene were higher in the color pool, while others were lower in the white pool. Thus, there was no clear pattern to suggest that differences in carotenoid synthesis gene expression were involved in differences in fruit flesh color.

**Table 3.** Differentially expressed genes of the carotenoid synthesis pathway.

| **<u>GeneID</u>** | **<u>Description</u>** | **<u>logFC<sup>1</sup></u>** | **<u>Position<sup>2</sup></u>** |
| --- | --- | --- | --- |
| CsGy4G007210.1 | Phytoene synthase 3 (PSY) | 0.66 | upstream |
| CsGy4G003270.1 | 15-cis-zeta-carotene isomerase (ZISO) | -0.79 | upstream |
| CsGy1G032330.1 | Zeta-carotene desaturase (ZDS) | -0.61 | upstream |
| CsGy3G006450.1 | prolycopene isomerase 1 (CRTISO) | 0.74 | upstream |
| CsGy4G000680.1 | Lycopene beta cyclase (b-LCY) | -0.86 | upstream |
| CsGy3G007100.1 | Cytochrome P450, putative (BCH) | 1.03 | downstream |
| CsGy3G017310.1 | Beta-carotene hydroxylase (BCH) (Ore gene) | -1.28 | downstream |
| CsGy2G014000.1 | Zeaxanthin epoxidase (ZEP) | -0.99 | downstream |
| CsGy2G008070.1 | violaxanthin de-epoxidase (VDE) | 0.74 | downstream |
| CsGy2G015170.1 | violaxanthin de-epoxidase (VDE) | 1.49 | downstream |
| CsGy4G006910.1 | Carotenoid cleavage dioxygenase 4 (CCD) | -2.03 | downstream |
| CsGy4G007460.1 | 9-cis-epoxycarotenoid dioxygenase (NCED) | -1.29 | downstream |
| CsGy6G035690.1 | 9-cis-epoxycarotenoid dioxygenase (NCED) | -2.88 | downstream |

<sup>1</sup>Positive logFC means higher in color pool, negative logFC means higher in white pool
<sup>2</sup>Position in carotenoid synthesis pathway relative to beta-carotene

Since a mutation in the Or gene of cauliflower, which encodes a DnaJ chaperone protein, leads to carotenoid accumulation, we also checked for differential expression of DnaJ chaperone genes in the color and white pools. Nine DnaJ chaperone genes had higher expression in the color pool. However, none of these transcripts corresponded to the CsGy6G025020 gene that is most homologous to the Or gene.

### Enriched SNPs in each pool

From the RNA sequencing reads, we were also able to identify SNP occurrences in the three color pool samples and the three white pool samples, relative to the reference genome (Table S3). Only one of the carotenoid synthesis transcripts had any SNPs; CsGy1G032330, which encodes ZDS, had three SNPs. However, these SNPs are unlikely to be involved in color accumulation, because although they were found mainly in the color pool, they were not homozygous. In addition to the carotenoid synthesis pathway, we checked for SNPs in DnaJ chaperone protein-encoding genes. Most interestingly, CsGy6G025020, which is homologous to the Or gene, had an indel SNP, an insertion of a G, at position 923. This indel was homozygous in all of the color pool samples. This insertion was not present in any of the white pool samples. In the deduced protein sequence from the reference and color pool transcripts, this insertion in the color pools would result in a translation frame shift followed by early termination of the protein (Fig. 3).

**Figure 3.**
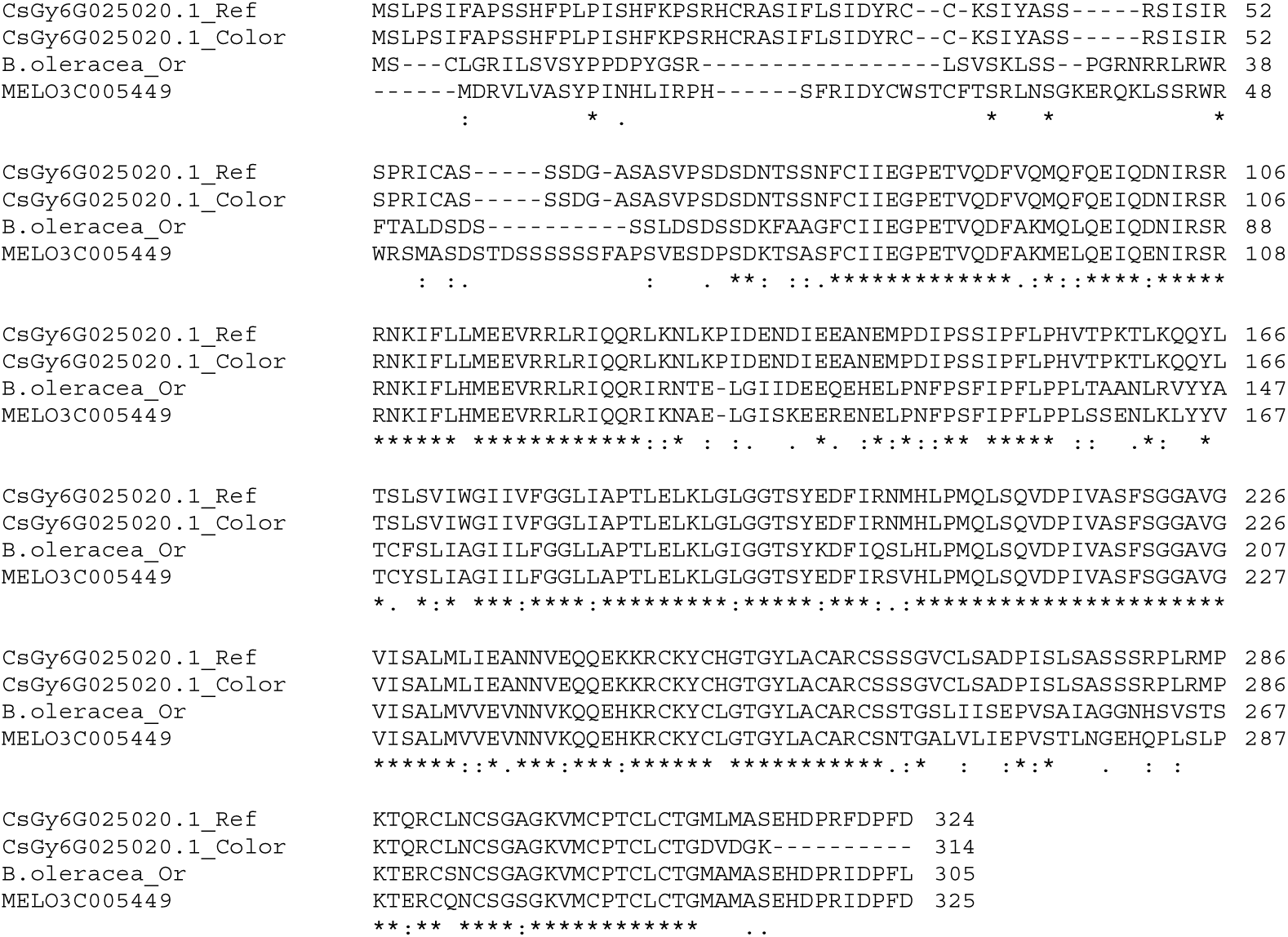
ClustalW alignment of deduced amino acid sequences of *Orange* (*Or*) genes from cucumber, cauliflower, and melon. CsGy6G025020.1_Ref, deduced sequence from reference transcriptome for *Or* homolog; CsGy6G025020.1_Color, deduced sequence from color pool with G indel at base 923. Frameshift occurs at AA 310; B.oleracea_*Or*, deduced sequence from wild-type *Or* gene of cauliflower; MELO3C005449, deduced sequence from melon *Or* gene.

### Carotenoid analysis in orange and white fruits

To determine whether the orange coloration of the cucumber flesh was due to ß-carotene, we used HPLC to determine ß-carotene levels in an orange fruit, a green fruit, and a white fruit (Fig. 2B). The white fruit had only a trace of ß-carotene, while the orange fruit had much greater quantities, and the green fruit had an intermediate level (Fig. 4). In addition to the ß-carotene peak, the green and orange fruits had several unidentified peaks that were eluted at earlier retention times. Thus, the colored fruits contained higher quantities of ß-carotene than the white fruits, and also more of some unknown compounds.

**Figure 4.**
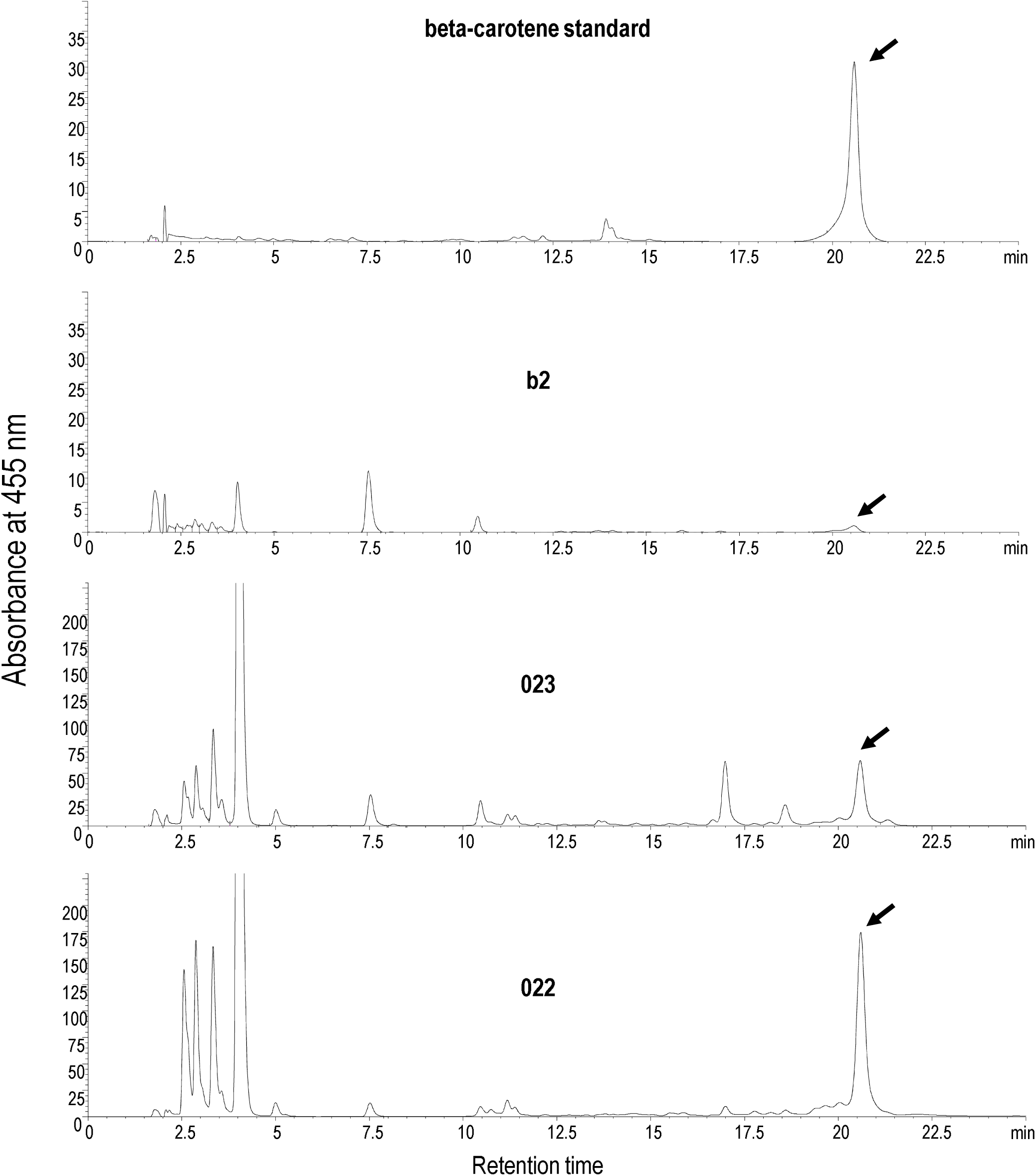
HPLC output for carotenoid analysis. Upper trace, beta-carotene standard; b2 trace, white-fleshed cucumber sample; 023 trace, green-fleshed cucumber sample; 022 trace, orange-fleshed cucumber sample. See Fig. 2B for photograph of white, green, and orange cucumbers.

## Discussion

### Previous genetic basis for orange cucumber

Our overall goal in this project was to identify genetic factors in cucumber that are responsible for orange fruit coloration. We confirmed that the orange coloration is due, at least in part, to ß-carotene (Fig. 4). In some instances, expression of *PSY* has been identified as a limiting factor in ß-carotene synthesis (Cazzonelli and Pogson, 2010). However, *PSY* expression and that of other genes in the carotenoid synthesis pathway do not seem to be likely causes of orange coloration, since there was no clear pattern of differential expression in either the color or white pools, upstream or downstream of ß-carotene (Table 3). In the closely related species *Cucumis melo* (melon), F_3_ families from a population with green-fleshed or orange-fleshed fruits were pooled by developmental time and fruit color (green or orange) (Chayut *et al.*, 2015). Despite the differences in ß-carotene concentration, expression of carotenoid synthesis genes was similar between the green and orange pools, consistent with our results.

SNPs in carotenoid synthesis genes were not detected in this study, except for some in a ZDS gene, which did not strictly segregate with the pools. In the previously studied orange cucumber fruits of the XIS lines, a mutation in *BCH1* was responsible for accumulation of ß-carotene in fruits (Qi *et al.*, 2013). However, we did not detect any SNPs in this gene in the cucumber lines in this study. Therefore, it is unlikely that changes in expression or function of ß-carotene synthesis or degradation genes underlie the orange coloration in fruits of these populations.

### The *Or* gene is mutated in the color pool

In cauliflower, the *BoOr* mutation in a DnaJ chaperone protein led to formation of chromoplasts and accumulation of ß-carotene in florets (Lu *et al.*, 2006). This cauliflower mutation resulted from a transposon. Similarly, in melon, a SNP in the *Or* gene resulted in an arginine to histidine substitution (Tzuri *et al.*, 2015), which segregated perfectly with orange or green fleshed fruits (Chayut *et al.*, 2015). In our study, we did detect an indel SNP in the homolog of the *BoOr* or *CmOr* genes, which was an extra G that resulted in a frame-shift and premature stop codon (Fig. 3). This mutation was homozygous all three samples in the orange pool. Thus, we believe it is likely to be necessary for orange coloration in these lines. The SNP in our cucumber samples occurred later in the translated protein than either the cauliflower or melon mutations. Thus, it seems that the *Or* gene may be quite sensitive to mutations, several of which at different positions (Lu *et al.*, 2006, Tzuri *et al.*, 2015) are enough to allow accumulation of ß-carotene inside chromoplasts that develop due to the mutation.

### Additional genetic factors

Although our study indicates a genetic basis for orange cucumber that differs from that of the XIS cucumber lines, larger population sizes and/or advanced generations could be studied using the *Or* SNP as a genetic marker to calculate genetic ratios. Although the *Or* SNP was homozygous in the color pool, with larger populations, larger pools composed of only orange fruits could be made, and additional pools with green or yellow flesh could allow a finer-scale dissection of gene expression and SNP differences that may be associated with flesh color. The PI 200815 line, which has yellow flesh and is one of the parents used in this study, was studied previously. One study attributed the yellow coloration to two recessive genes called v and *w* (Kooistra 1971), while a more recent study attributed the yellow coloration to a single recessive gene called *yf* (Lu *et al.*, 2015). The current study design did not allow us to accurately quantify color and segregation ratios, but there were fruits with varying degrees of color (Table 1) and full orange fruits were rare (2 of 126), suggesting that polygenic inheritance is likely, and that the *Or* SNP is not the only factor. Although this study did not allow us to identify the *yf* genes or other unknown genes, it seems clear that the *BCH1* gene is not the key factor in these populations, since there were no SNPs in it.

It might also be useful to study these populations separately, rather than pooling them. Because of small fruit numbers, our pools had uneven contributions of each parent in pools (Table 2). This pooling imbalance could have led us to miss other important DEGs or SNPs that might be more apparent in biparental populations. Based on our sampling of each population, there may be different important genes segregating in each one. For example, the PI 200815 X PI 163217 population had predominantly white-fleshed fruits, while the PI 606066 × 163217 population had green+yellow flesh as the most frequent coloration, and the PI 606066 X PI 200815 had a large number of green-fleshed fruits (Table 1, Table S1). Thus, studying each population separately might yield additional information about the genetic basis of green, yellow, or orange flesh in cucumber fruits.

## Conclusions

This study indicates that there are other sources of genetic variation for flesh color in cucumber, including the *Or* gene, and that it is not limited to the *BCH1* gene that is responsible for orange flesh coloration in the XIS populations. This new information gives plant breeders additional germplasm to use for breeding ß-carotene-enhanced cucumbers. It should be possible to use multiple sources of ß-carotene accumulation (e.g. the *BCH1* mutant from XIS, *Or* mutant lines from these PI lines, and other unidentified genes) to develop new varieties with improved nutritional value and a novel appearance that might be popular with consumers. Additionally, our dataset could be useful for datamining for future studies.

## Acknowledgements

This work was partially funded by a grant to BMW from the University of Nebraska Fleming Horticulture Research Fund. The authors thank Sam Polk for assistance with growing populations and sampling the fruits. The authors thank Edgar Cahoon for the use of the HPLC. The UNMC DNA Sequencing Core Facility receives partial support from the Nebraska Research Network In Functional Genomics NE-INBRE P20GM103427-14, The Molecular Biology of Neurosensory Systems CoBRE P30GM110768, The Fred & Pamela Buffett Cancer Center - P30CA036727, The Center for Root and Rhizobiome Innovation (CRRI) 36-5150-2085-20, and the Nebraska Research Initiative.

